# Age-dependent changes in transcription factor FOXO targeting in *Drosophila melanogaster*

**DOI:** 10.1101/456426

**Authors:** Allison Birnbaum, Xiaofen Wu, Marc Tatar, Nan Liu, Hua Bai

## Abstract

FOXO transcription factors have long been associated with longevity control and tissue homeostasis. Although the transcriptional regulation of FOXO have been previously characterized (especially in long-lived insulin mutants and under stress conditions), how normal aging impacts the transcriptional activity of FOXO is poorly understood. Here, we conducted a chromatin immunoprecipitation sequencing (ChIP-Seq) analysis in both young and old wild-type fruit flies, *Drosophila melanogaster*, to evaluate the dynamics of FOXO gene targeting during aging. Intriguingly, the number of FOXO-bound genes dramatically decreases with age (from 2617 to 224). Consistent to the reduction of FOXO binding activity, many genes targeted by FOXO in young flies are transcriptionally altered with age, either up-regulated (FOXO-repressing genes) or down-regulated (FOXO-activating genes). In addition, we show that many FOXO-bound genes in wild-type flies are unique from those in insulin receptor substrate *chico* mutants. Distinct from *chico* mutants, FOXO targets specific cellular processes (e.g., actin cytoskeleton) and signaling pathways (e.g., Hippo, MAPK) in young wild-type flies. FOXO targeting on these pathways decreases with age. Interestingly, FOXO targets in old flies are enriched in cellular processes like chromatin organization and nucleosome assembly. Furthermore, FOXO binding to core histone genes is well maintained at aged flies. Together, our findings provide new insights into dynamic FOXO targeting under normal aging and highlight the diverse and understudied regulatory mechanisms for FOXO transcriptional activity.

## Introduction

The process of aging is accompanied by a decline in physiological function and cellular maintenance. It is known that aging dramatically alters gene expression and transcription factor activity (Lopez-Otin, Blasco, Partridge, Serrano, & Kroemer, 2013). The protein family of Forkhead Box subfamily O transcription factors, or FOXO, has been shown to play an important role in growth, development, stress resistance, and longevity (Greer & Brunet, 2005). FOXO functions downstream of insulin/insulin-like growth factor (insulin/IGF) signaling and is negatively regulated by PI3K-Akt pathway (Brunet et al., 1999). FOXO transcriptionally regulates numerous target genes involving metabolism, cell cycle progression, stress, and apoptosis (Kitamura et al., 2002; Kops et al., 2002; Martins, Lithgow, & Link, 2016; Medema, Kops, Bos, & Burgering, 2000). Additionally, FOXO proteins were first implemented in lifespan extension in *Caenorhabditis elegans* where insulin-like receptor mutant *daf-2* extends lifespan via FOXO homolog *daf-16* (Kenyon, Chang, Gensch, Rudner, & Tabtiang, 1993). This lifespan extension through insulin/IGF signaling has been observed across species, from worm to fly to mammal (Holzenberger et al., 2003; Kenyon et al., 1993; Tatar et al., 2001). Studies have found that lifespan extension effects of insulin/IGF deficiency depend on FOXO activity, probably through the transcriptional regulation of key longevity assurance pathways such as xenobiotic resistance (Slack, Giannakou, Foley, Goss, & Partridge, 2011; Yamamoto & Tatar, 2011). However, how FOXO elicits this response remains to be fully elucidated.

FOXO activity is not solely dependent on insulin/IGF signaling. FOXO proteins undergo posttranslational modifications in response to other cellular stress signals. Oxidative stress promotes Jun-N-terminal Kinase (JNK)-dependent phosphorylation of mammalian FOXO4 and its nuclear translocation. FOXO proteins can also be activated and phosphorylated by mammalian Sterile 20-like kinase 1 (MST1), to extend lifespan (Dansen & Burgering, 2008; Essers et al., 2004; Lehtinen et al., 2006). In response to DNA damage, cyclin-dependent kinase 2 (CDK2) can phosphorylate and regulate mammalian FOXO1 to delay cell cycle progression and induce apoptosis (Huang & Tindall, 2006). FOXO proteins are also involved in tumor suppression activity and responds to oncogenic stress (Dansen & Burgering, 2008). Interestingly, two recent chromatin immunoprecipitation-sequencing (ChIP-Seq) studies revealed that FOXO proteins are enriched at the promoters of many target genes in well-fed wild-type *C. elegans* and *Drosophila* (Alic et al., 2011; Riedel et al., 2013).

Although insulin/IGF signaling is well-known aging regulators, how insulin/IGF signaling is altered during normal aging remains largely unclear. It is generally believed that insulin/IGF signaling declines with age. This is primarily based on age-dependent decrease in the expression of FOXO target genes (Demontis & Perrimon, 2010; Rera, Clark, & Walker, 2012). However, it remains to be determined how aging impacts FOXO transcriptional activity and DNA binding capacity of FOXO transcription factors. Here, we conducted a ChIP-Seq analysis to investigate FOXO binding dynamics under normal aging in *Drosophila*. Intriguingly, we found that the number of FOXO-bound regions sharply decrease with age. The age-related decrease in FOXO binding is correlated with either the transcriptional activation of FOXO-repressing genes, or the downregulation of FOXO-activating genes during normal aging. Furthermore, we observed strong FOXO nuclear localization in well-fed wild-type flies, while FOXO targets distinct sets of genes between wild-type and insulin mutants. Taken together, our findings provide new evidence linking age-dependent FOXO transcriptional activity to its role in longevity control and tissue maintenance.

## Results

### FOXO exhibits constitutive nuclear localization in young and old adult fat body

To examine whether *Drosophila* FOXO activity changes with aging, we first performed immunofluorescent staining using a polyclonal antibody against *Drosophila* FOXO to monitor the FOXO nuclear localization in wild-type flies (*yw^R^*) at two different ages, two-week-old (young flies) and five-week-old (aged flies). Intriguingly, FOXO proteins exhibited constitutive nuclear localization in abdominal fat body tissue of well-fed wild-type female flies (*yw^R^*), where insulin/IGF signaling is presumably active (Figure 1A). The constitutive nuclear localization of FOXO was also found in another wild-type line, *Oregon R (OreR)* (Figure S1). FOXO proteins remained nuclear localization during aging, while the colocalization of FOXO with nuclear DAPI staining slightly declined in aged fat body tissue (Figure 1A-1B). Compared to adult fat body, indirect flight muscles from both two-week-old and five-week-old female flies showed low FOXO nuclear localization (Figure 1C, 1D, S1). Thus, these results suggest that FOXO could be activated in well-fed wild-type flies to regulate the expression of its target genes, which is consistent with recent ChIP-Seq studies (Alic et al., 2011; Riedel et al., 2013).

**Figure 1.**
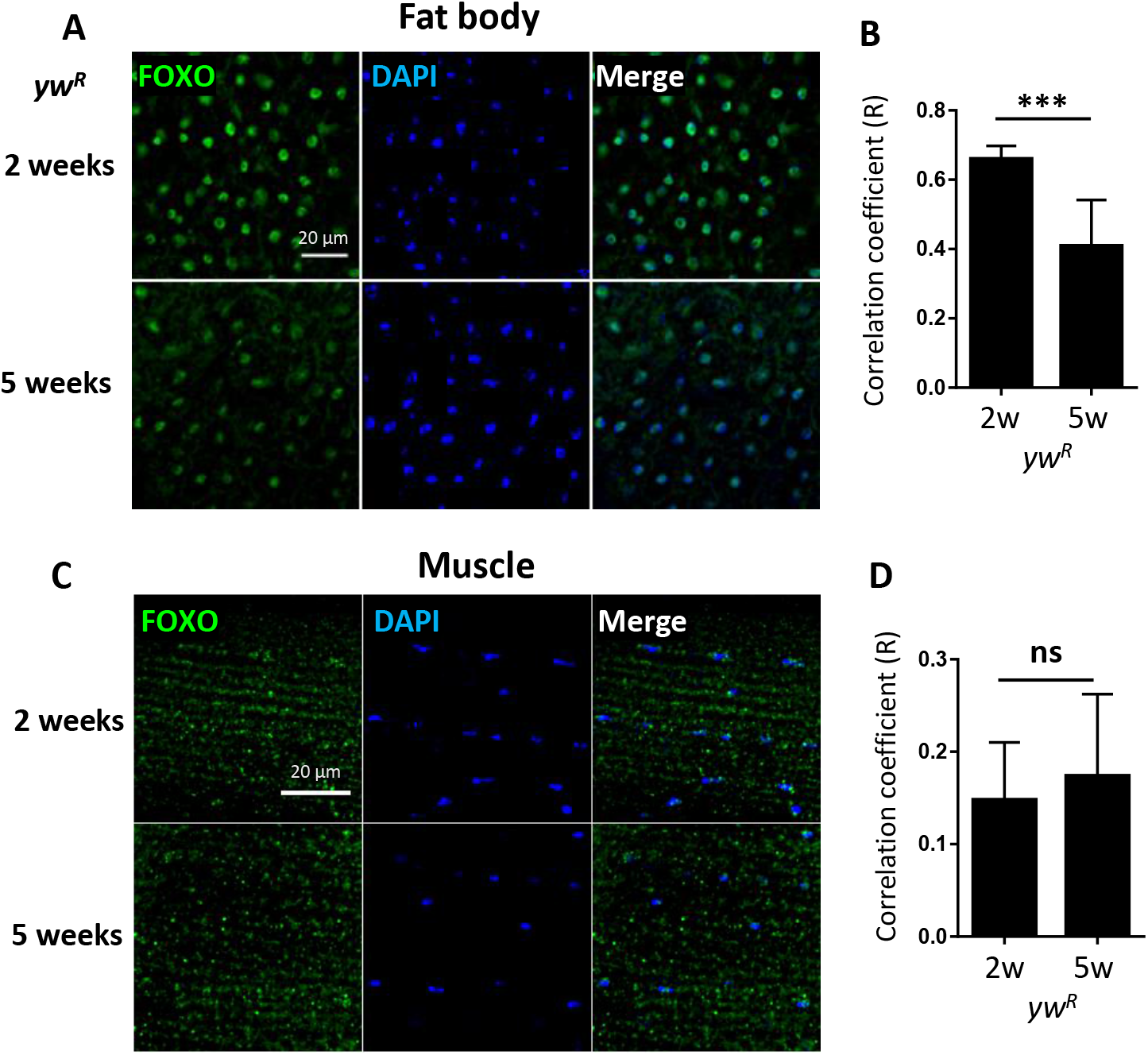
FOXO exhibits constitutive nuclear localization in young and old adult fat body. **A)** Abdominal fat body of wild-type flies (*yw^R^*) stained with anti-FOXO at young (2 weeks) and old ages (5 weeks). **B)** Quantification of Pearson correlation coefficient (R) between FOXO and DAPI staining in fat body tissue. **C)** FOXO immunostaining in young and old indirect flight muscles of wild-type flies (*yw^R^*). **D)** Quantification of Pearson correlation coefficient (R) between FOXO and DAPI in indirect flight muscles. Scale bar: 20 μm. Student *t*-test (***, *p*< 0.001; ns: not significant).

### ChIP-Seq analysis reveals age-dependent reduction of FOXO-targeted DNA binding

To further investigate the FOXO transcriptional activity under normal aging, we performed ChIP-Seq analysis on young (2-week) and aged (5-week) female wild-type flies. Using Illumina high-throughput sequencing, we obtained a total of 261 million reads from FOXO ChIP and input DNA samples at two ages. On average, 90.08% of unique reads were mapped to annotated *Drosophila* reference genome (Figure S2A, Table S1:List 24). Intriguingly, our ChIP-Seq analysis revealed that the number of FOXO-bound genomic regions (based on MACS2 peak calling) dramatically decreased with age (Figure 2A). There were 9273 peaks identified in young flies (corresponding to 2617 protein coding genes), whereas in aged flies only 1220 peaks (224 genes) were detected (Figure 2A, Table S1:List 5-8). About 170 genes were shared between two ages. For most of the peaks, a reduction in peak size or a disappearance of peaks was observed in aged flies (Figure 2B), while the FOXO binding to a few genomic regions remained unchanged during aging (Figure 2C). The reduction of FOXO-bound regions was not due to the decreased quantity of immunoprecipitated genomic DNA (data not shown). In fact, equal amount of ChIP and input DNA samples were used to generate Illumina sequencing libraries. In addition, a correlation matrix plot showed that the reads from 2-week-old FOXO ChIP samples were most divergent from the input and 5-week-old ChIP samples, further suggesting the differential FOXO-DNA binding activity between young and aged flies (Figure S2B).

**Figure 2.**
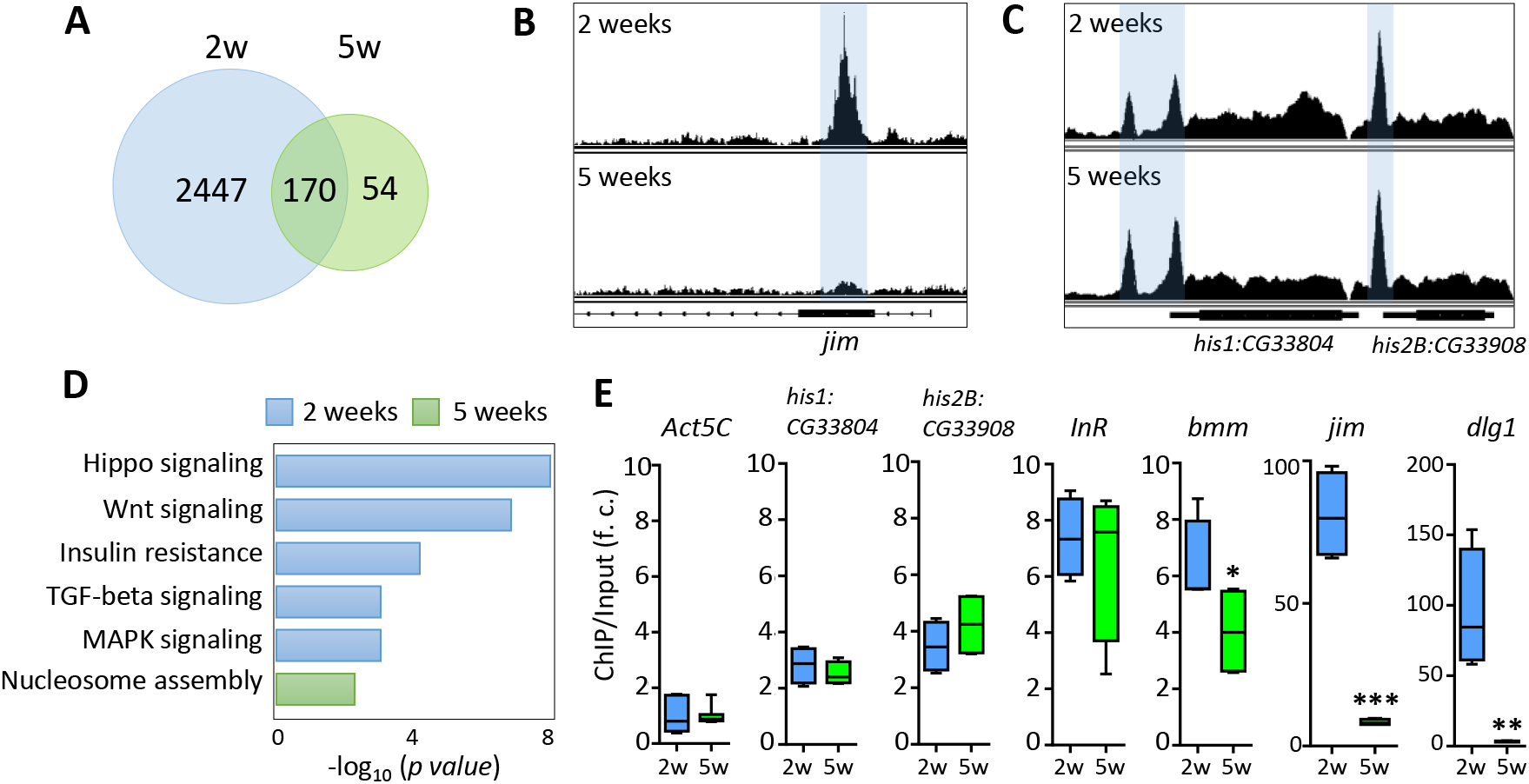
FOXO binding activity decreases with age. **A)** The number of genes targeted by FOXO at young (2 weeks) and old ages (5 weeks). **B)** Age-dependent FOXO binding at *jim* locus. **C)** Age-dependent FOXO binding at *his1:CG33804* and *his2B:CG33908* loci. **D)** GO terms for FOXO-targeted pathways uniquely enriched in young or old flies. **E)** qPCR validation of the FOXO binding enrichment at the selected FOXO targeted genomic loci. FOXO binding at Act5C locus serves as an internal control. The enrichment value is calculated as the fold-change (f. c.) of the FOXO binding (ChIP vs. Input) between FOXO-targeted loci and Act5C locus. Student *t*-test (***, *p*<0.001; **, *p*<0.01; *, *p*<0.05).

Pathway analysis revealed that FOXO target genes at young ages were enriched in pathways like Hippo, Wnt, TGF-beta, MAPK, and insulin resistance pathways (Figure 2D, Table S1:List 10). FOXO was also targeting genes involved in nervous system development, motor neuron stabilization, and regulation of synaptic tissue communication (Table S1:List 10). Additionally, we found that FOXO bound to the genomic regions containing key autophagy regulators (*Atg3, Atg17, Tor, wdb, Pten*), which is consistent to previous known functions of FOXO in autophagy and tissue homeostasis (Demontis & Perrimon, 2010). Many Rho and small GTPase proteins, as well as actin cytoskeleton pathways, are also targeted by FOXO at young ages. Many of these FOXO-targeted pathways were absent in aged flies. Instead, processes like nucleosome assembly and chromatin organization were enriched as FOXO-bound targets in aged flies (Figure 2D, Table S1:List 11). Interestingly, strong FOXO binding was maintained at many core histone genes at old ages (Figure 2C, 2E).

The age-dependent changes in FOXO binding activity were verified by quantitative PCR (ChIP-qPCR). The FOXO binding to the promoters of two known target genes, insulin receptor *InR* and adipose triglyceride lipase *bmm*, were first tested in ChIP-qPCR analysis (Figure 2E). FOXO showed similar binding enrichment (6~7 fold) at *InR* locus between young and old ages (Figure 2E). On the other hand, the FOXO binding to *bmm* promoter slightly decreased with age (Figure 2E). We also confirmed that FOXO binding remained unchanged at two histone loci (*his1:CG33804* and *his2B:CG33908*), while the FOXO enrichment at two newly identified target genes, *jim* (C2H2 zinc finger transcription factor) and *dlg1* (a key factor for the formation of septate junctions and synaptic junctions), decreased dramatically at old ages (from 80~90-fold to 3~8-fold) (Figure 2E). Thus, our ChIP-qPCR analysis confirmed that FOXO binding activity was altered in many target loci during normal aging.

### FOXO-bound genes show age-dependent transcriptional changes

We next examined whether age-dependent changes in FOXO binding is correlated to age-regulated transcription of FOXO target genes. To do so, we first compared our FOXO ChIP-Seq results to previously published aging transcriptomic analysis on aging *Drosophila* tissues, such as fat body and head tissue. Out of 2447 FOXO target genes (uniquely bound by FOXO at young ages), 408 of them were differentially expressed in aging fat body (172 downregulated, 236 upregulated) (Figure 3A, Table S1:List 12), while 845 target genes were differentially expressed in aging head tissue (626 downregulated, 219 upregulated) (Figure 3C, Table S1:List 13). Interestingly, a majority of the FOXO-bound genes showed no age-related transcriptional changes, which is similar to previous studies showing the FOXO binding at the promoters of large number of so-called poised genes (Webb, Kundaje, & Brunet, 2016; Webb et al., 2013). Gene ontology analysis revealed that FOXO target genes differentially expressed in aging fat body were enriched for processes and signaling pathways like chromatin organization, histone modification, hippo signaling, peroxisome, and hormone biosynthesis (Figure 3B, Table S1:List14). On the other hand, the differentially expressed FOXO targets in aging head tissue were enriched for pathways and processes involving Wnt, Hippo, G protein-couple receptor (GPCR), axon guidance, synapse organization, and actin cytoskeleton (Figure 3D, Table S1:List15).

**Figure 3.**
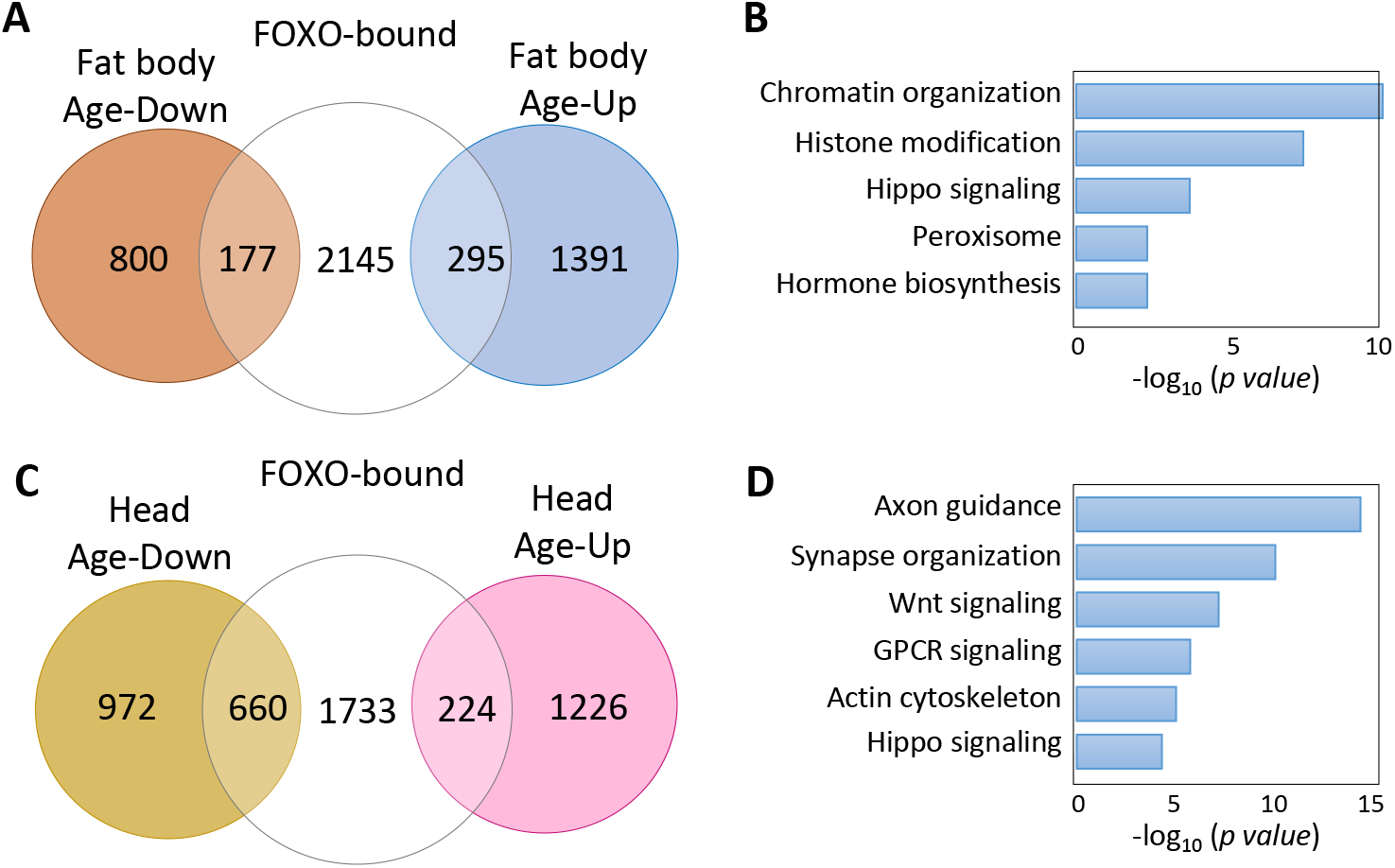
FOXO target genes show age-dependent transcriptional changes. **A)** The number of FOXO-bound genes that are differentially expressed in aging fat body. **B)** Representative biological processes enriched for age-regulated FOXO targets in fat body. **C)** The number of FOXO-bound genes that are differentially expressed in aging head tissue. **D)** Representative biological processes enriched for age-regulated FOXO targets in adult head tissue.

Although many FOXO-bound target genes exhibited differential expression during aging, it remains unclear whether decreased FOXO-binding activity at old ages contributes to age-dependent transcriptional changes of these FOXO target genes. To further determine the relationship between FOXO binding and transcriptional changes of FOXO target genes, we performed a RNA-Seq analysis using head tissues dissected from wild-type flies and a *foxo* null mutants (*foxo^c431^*), a site-specific deletion mutant generated by CRISPR/Cas9 (Figure 4A–4B). Out of 2617 FOXO-bound target genes, 101 of them were upregulated in *foxo^c431^* mutants, while 300 were downregulated in the mutants (Figure 4C, Table S1:List 16), suggesting that FOXO binding might be important to repress or activate at least a subset of target genes. Based on these data, FOXO target genes can be sorted into three classes, FOXO-repressing (101 genes), FOXO-activating (300 genes), and FOXO-no regulation (1621 genes).

**Figure 4.**
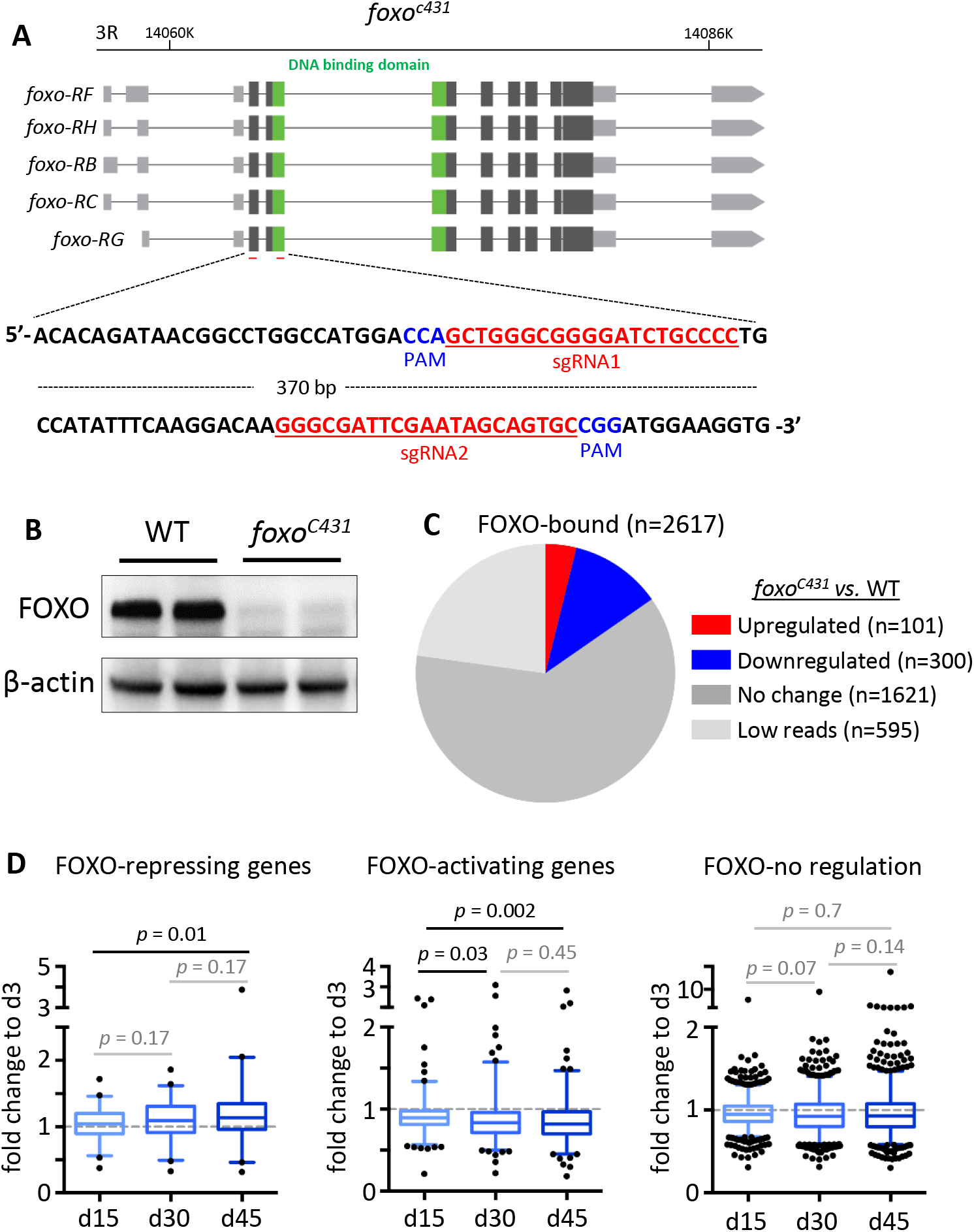
The altered of FOXO binding correlates with age-related transcriptional changes of FOXO targets. **A)** The diagram showing *foxo* locus and the target sites of the guiding RNAs (highlighted in red) used to generate *foxo^c431^* loss-of-function mutants by CRISPR/Cas9 mutagenesis. PAM: Protospacer adjacent motifs (highlighted in blue). **B)** Western blots to verify the expression of FOXO proteins in *foxo^c431^* loss-of-function mutants. β-actin as a loading control. **C)** The number of FOXO target genes that are differentially expressed between *foxo^c431^* mutants and wild-type flies. **D)** Age-dependent transcriptional changes of FOXO target genes. Boxplots represent the mean fold change of genes at Day 15 (d15), Day 30 (d30) and Day 45 (d45), relative to that of Day 3 (d3) in aging head tissue (Student *t*-test).

We next asked how reduced FOXO binding during aging impacts the expression of FOXO target genes. To do this, we first constructed new transcriptomic profiles from wild-type head tissue at four different ages, 3d, 15d, 30d, and 45d (Table S1:List 17). Interestingly, among three classes of FOXO target genes, FOXO-repressing genes exhibited an increased expression in old flies, whereas FOXO-activating genes were progressively downregulated with age. Expression of FOXO-no regulation genes, on the other hand, did not significantly change during aging (Figure 4D). Taken together, these results suggest that age-associated decrease in FOXO binding might contribute directly to the transcriptional alterations of FOXO target genes in old flies.

### FOXO binding differs between wild-type and insulin/IGF mutants

FOXO binding activity has been primarily studied by evaluating its response to IIS signaling (Alic et al., 2011; Bai, Kang, Hernandez, & Tatar, 2013; Murphy, 2006; Riedel et al., 2013; Webb et al., 2016). However, our observations on FOXO nuclear localization and DNA binding in well-fed wild-type flies suggest that there might be distinct FOXO transcriptional activity independent of insulin/IGF signaling. To test this possibility, we compared FOXO ChIP-Seq datasets from the present study (young wild-type) and our previous analysis on insulin receptor substrate *chico* mutants (Bai et al., 2013). Intriguingly, large number of FOXO-bound genes were not shared between wild-type and *chico* mutants. There were1992 FOXO target genes unique to wild-type, while 1393 genes unique to *chico* mutants (Figure 5A, Table S1:List 18). Furthermore, distinct FOXO targets between wild-type and *chico* mutants were differentially expressed with age (Figure 5B). About 844 age-regulated genes were only bound by FOXO in wild-type flies, while 577 genes unique to *chico* mutants (Table S1:List 19). We found that age-regulated FOXO targets unique to *chico* mutants were enriched in metabolic pathway and oxidative-reduction, while those unique to wild-type flies were enriched for chromatin organization, axon guidance, Hippo and MAPK signaling pathways (Figure 5C, Table S1:List 20-21). When examining each pathway in detail, we noticed that FOXO targets in Hippo and MAPK/EGFR signaling pathways were found in both wild-type and *chico* mutants, although different target genes were apparent between the two conditions (Figure S3–S4).

**Figure 5.**
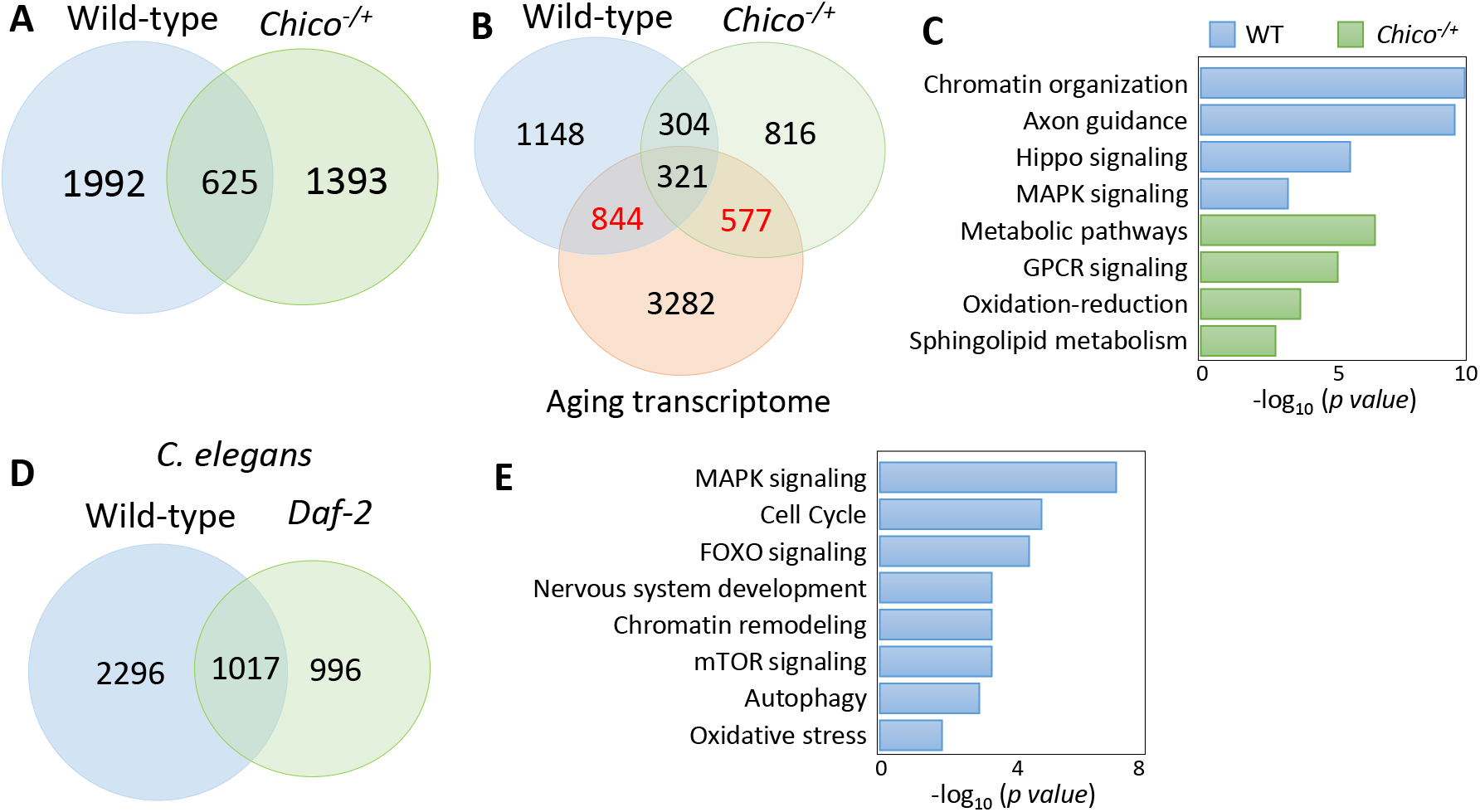
FOXO binding differs between wild-type and insulin/IGF mutants. **A)** Comparison of FOXO target genes between wild-type and *chico* mutants. **B)** Overlap between age-dependent differentially expressed genes (fat body and head) and FOXO-bound targets (wild-type and *chico* mutants). **C)** GO terms uniquely enriched in wild-type or *chico* mutants. **D)** Daf-16-bound targets genes in wild-type *C. elegans* and *Daf-2* mutants. **E)** Shared pathways targeted by both fly FOXO and worm Daf-16 in wild-type animals. Enriched *C. elegans* GO terms are shown.

To test if distinct FOXO binding activity observed between wild-type flies and insulin/IGF mutants is conserved across species, we reanalyzed the recent *C. elegans Daf-16* ChIP-seq study (Riedel et al., 2013). Interestingly, wild-type worms also showed different Daf-16 binding activity from *daf-2* mutants. There were 2296 genes uniquely bound by Daf-16 to wild-type worms, while 996 were unique to *daf-2* mutants (Figure 5D, Table S1:List 22). Gene ontology analysis showed that FOXO transcription factors targeted similar pathways in wild-type flies and worms. These pathways were MAPK signaling, cell cycle, FOXO signaling, nervous system development, chromatin remodeling, mTOR signaling, autophagy, and oxidative stress (Figure 5E, Table S1:List 23). Thus, insulin/IGF-independent FOXO transcriptional activity may be an evolutionarily conserved cellular mechanism.

### Enriched FOXO motifs in wild-type flies

A signature of FOXO targeting is the 8-nucleotide long canonical binding motif, 5’-TTGTTTAC-3’, which is conserved across species (Bai et al., 2013; Furuyama, Nakazawa, Nakano, & Mori, 2000; Webb et al., 2013). This motif is typically found upstream of the gene coding site in the enhancer or promoter region (Eijkelenboom, Mokry, Smits, Nieuwenhuis, & Burgering, 2013; Webb et al., 2013). To search for FOXO consensus sequence in the FOXO-bound genomic regions found in young wild-type flies, we conducted motif analysis using the Homer motif finding tool. We used peaks with at least a 2-fold enrichment that were less than 2000 bp in length, and we searched for motifs within 200 bp surrounding the peak region. When insect motif databases were used, we identified only one known motif for Trl (p < 10^-70^), a GAGA-factor that also found in previous ChIP-Seq data from *C. elegans* (Riedel et al., 2013) (Table 1). When searching against known mammalian motifs, a motif for FOXO1 (with canonical consensus, TGTTTAC) was detected with low significance (p < 10^-4^). (Table 1). Next, using *de novo* motif search we found that FOXO-bound regions were enriched with motifs for transcription factors hb, Adf1, and Aef1. Lastly, we performed Homer *de novo* motif search and identified a motif for RAP1, a *Saccharomyces cerevisiae* gene that is part of the Myb/SAINT domain family, which was also found in a previous *Drosophila* FOXO ChIP-on-ChIP study (Alic et al., 2011). Together, these findings suggest that in wild-type flies FOXO may recognize a unique set of motifs that is different from the canonical consensus sequence.

**Table 1:**
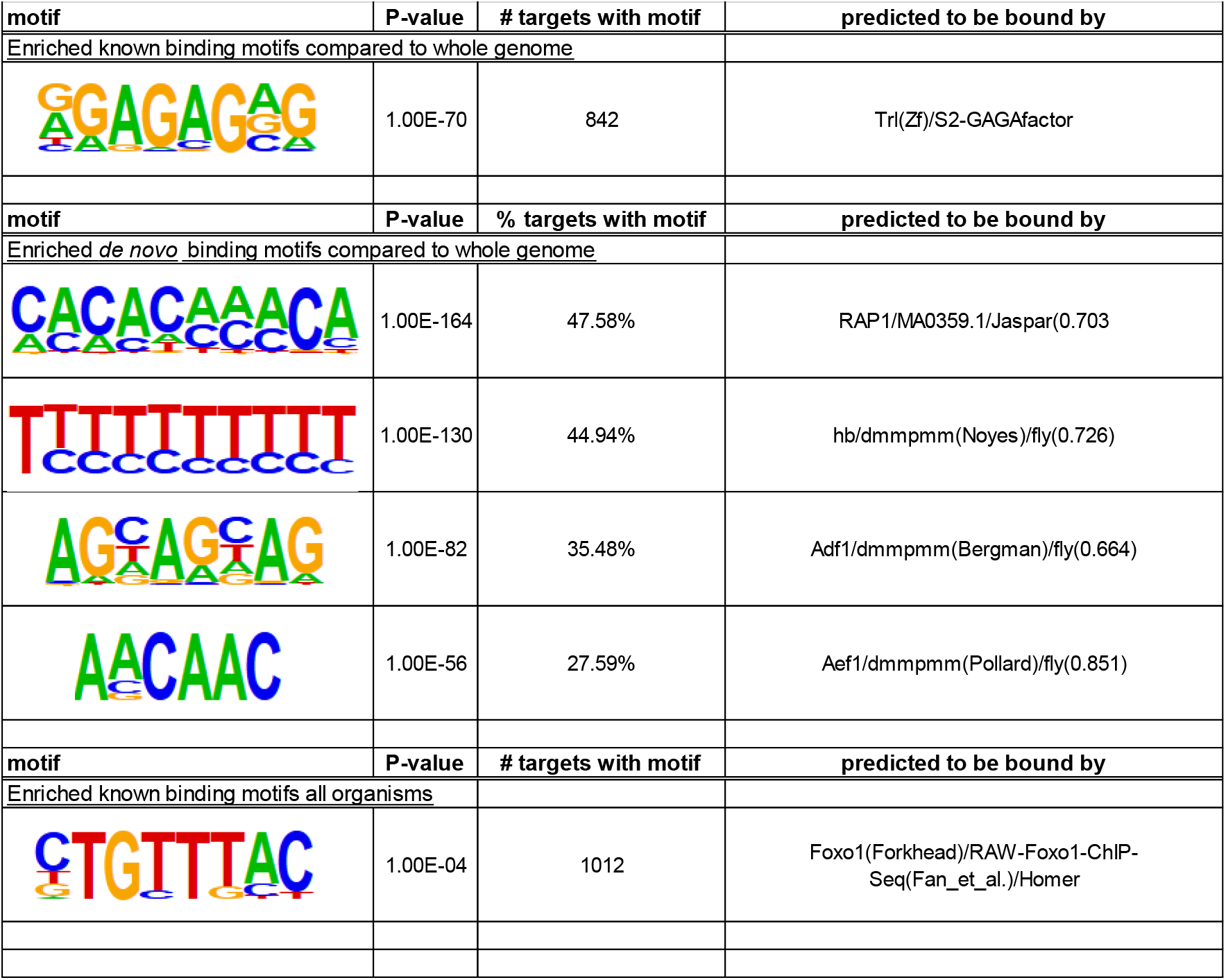
Lists of motifs that are enriched among FOXO target sites in wild-type flies.

| motif | P-value | # targets with motif | predicted to be bound by |
| --- | --- | --- | --- |
| Enriched known binding motifs compared to whole genome |  |  |  |
|  | 1.00E-70 | 842 | Trl(Zf)/S2-GAGAFactor |
| motif | P-value | % targets with motif | predicted to be bound by |
| Enriched *de novo* binding motifs compared to whole genome |  |  |  |
|  | 1.00E-164 | 47.58% | RAP1/MA0359.1/Jaspar(0.703 |
|  | 1.00E-130 | 44.94% | hb/dmmpmm(Noyes)/fly(0.726) |
|  | 1.00E-82 | 35.48% | Adf1/dmmpmm(Bergman)/fly(0.664) |
|  | 1.00E-56 | 27.59% | Aef1/dmmpmm(Pollard)/fly(0.851) |
| motif | P-value | # targets with motif | predicted to be bound by |
| Enriched known binding motifs all organisms |  |  |  |
|  | 1.00E-04 | 1012 | Foxo1(Forkhead)/RAW-Foxo1-ChIP-Seq(Fan_et_al.)/Homer |

## Discussion

As a key player in longevity control, FOXO transcription factors and their direct targets have been well characterized in many model systems (Alic et al., 2011; Bai et al., 2013; Riedel et al., 2013; Webb et al., 2013). However, whether and how FOXO transcriptional activity changes with age is unclear. In the present study, we performed a ChIP-Seq analysis to examine the FOXO binding activity during *Drosophila* aging. Intriguingly, genome-wide FOXO-binding underwent an immense reduction at old ages. Consistently, genes that are negatively regulated by FOXO showed an increased expression with age, whereas the FOXO-activating genes were downregulated in aged flies. Thus, age-associated decrease in FOXO binding is tightly linked to the transcriptional alterations of FOXO target genes at old ages. In addition, we found that FOXO targets distinct sets of genes between wild-type and insulin/IGF mutants across species, suggesting a conserved insulin/IGF-independent transcriptional regulation by FOXO transcription factors.

Changes in transcription factor binding patterns at different stages of life are not exclusive to FOXO. In *C. elegans*, FoxA/PHA-4 exhibits differential binding patterns at different stages of development to regulate organogenesis (Zhong et al., 2010). Similar to FOXO binding pattern, PHA-4 also exhibited binding at poised locations in the genome. The loss of specific FOXO targeting with age observed in the present study could be caused by either altered post-translational modification of FOXO, or changes in co-transcriptional regulation between FOXO and its partners. It is known that FOXO co-factors play an important role in fine-tuning FOXO transcriptional activity (Alic et al., 2011; Essers et al., 2004; Riedel et al., 2013; Webb et al., 2016). These co-factors include post-translational modifiers and nuclear interacting partners which aid FOXO in recruitment to target binding sites (Daitoku, Sakamaki, & Fukamizu, 2011; van der Vos & Coffer, 2008). A previous meta-analysis identified the binding motifs of many of novel transcription factors (EST, NRF and GATA factors) are enriched at FOXO target genes with age-related expression patterns (Webb et al., 2016), which suggests that the interplay between FOXO and these transcription factors may contribute to the altered FOXO transcriptional activity during normal aging. Certain mammalian FOXO co-factors, such as peroxisome proliferator-activated receptor gamma (PPARγ), and its coactivator (PGC-1α) interact with FOXO and compete for binding with FOXO and β-Catenin (Olmos et al., 2009; Polvani, Tarocchi, & Galli, 2012). FOXO acts as a repressor of PPARγ gene transcription, and this repression is lost later in life, suggesting a reduction of FOXO binding at PPARγ locus (Armoni et al., 2006; Polvani et al., 2012). Besides PPARγ and PGC-1α, many other transcription co-regulators and post-translational modifiers have been shown to be involved in transcriptional co-regulation of FOXO target genes, which may play important roles in modulating FOXO transcriptional activity during aging (Daitoku et al., 2011; van der Horst & Burgering, 2007).

Many FOXO-targeted cellular processes (e.g., nervous system development and actin cytoskeleton) and signaling pathways (e.g., Hippo, Wnt, TGF-beta, MAPK) are uniquely enriched in young wild-type, but not in *chico* mutants. Majority of these FOXO targets show age-dependent differential expression patterns. The Hippo pathway was initially characterized for its role in controlling organ size during development, but recently it has been shown to involve in autophagy, oxidative stress response, and aging (Lehtinen et al., 2006; Mao, Gao, Bai, & Yuan, 2015; Udan, Kango-Singh, Nolo, Tao, & Halder, 2003). In adult mice, suppression of Hippo signaling improved cell proliferation and heart tissue regeneration and is a regulator of tissue homeostasis (Heallen et al., 2013). Thus, Hippo signaling may be one of the major FOXO targets in the regulation of cellular homeostatic and longevity. MAPK signaling is involved in tissue homeostasis with aging (Jiang, Grenley, Bravo, Blumhagen, & Edgar, 2011; Lee & Sun, 2015), and is also enriched among FOXO-bound target genes in wild-type flies. Both the EGFR and JNK cascades of the MAPK signaling pathway are targeted by FOXO. The target genes involved in the EGFR signaling exhibit transcriptional alterations with age in the wild-type fly. In adult *Drosophila*, EGFR signaling is responsible for maintaining midgut epithelial homeostasis in the adult and has also been shown to regulate cytoskeletal modulation and autophagy (Hazan & Norton, 1998; Jiang et al., 2011; Tan, Lambert, Rapraeger, & Anderson, 2016). EGFR regulation of autophagy also impacts glial maintenance and degeneration of the nervous system (Lee & Sun, 2015). Our ChIP-Seq analysis places FOXO as an upstream regulator of MAPK/EGFR pathway to control autophagy and tissue maintenance during aging.

Our analysis also revealed that FOXO targets chromatin organization and nucleosome assembly processes. This finding suggests that FOXO may be involved in the maintenance of chromatin structure. Recent studies have shown that FOXO recruits SWI/SNF chromatin remodelers to specific target sites to regulate lifespan in *C. elegans* (Riedel et al., 2013). Changes in chromatin structure and overall loss of heterochromatin has long been an indicative measurement of aging (Larson & Yuan, 2012; Wood et al., 2010; Zhang et al., 2015). It is likely that FOXO plays an important role in maintaining chromatin structure and preventing age-related chromatin remodeling. Interestingly, we found that many core histone genes are targeted by FOXO. The binding of FOXO to these histone genes dramatically increases at old ages. It has been shown that the transcripts of histone genes increase during yeast replicative aging, but the levels of core histone proteins (e.g., H3, H2A) dramatically decrease with age (Feser et al., 2010). Overexpression of histones or mutation of histone information regulator (Hir) increase lifespan. How histone genes is transcriptional regulated during aging is unclear. Our findings suggest that FOXO might be one of the molecular mechanisms that contribute to altered histone expression during normal aging.

In summary, using a genome-wide approach we identified dynamic FOXO binding activity during *Drosophila* aging. Our findings further support the important role of FOXO in age-related transcriptional alterations and the regulation of tissue homeostasis and cellular maintenance pathways. Further investigation of the functional significance of the altered FOXO binding with age will be important in understanding how FOXO regulates organismal homeostasis and longevity.

## Experimental Procedures

### Fly culture and stocks

Flies were maintained at 25°C with 12 hour light/dark cycle, 60% humidity on agar-based diet with 0.8% cornmeal, 10% sugar, and 2.5% yeast. *yw^R^* flies (Bai et al., 2013) were used as wild-type for ChIP-Seq. *w^1118^* (Bloomington #5905) was used as a control genotype for *foxo* mutants in RNA-Seq analysis. Female flies were collected and sorted 1-2 days after eclosion. To age flies, vials contained 25-30 flies were transferred to fresh food every three days.

### CRISPR/Cas9 mutagenesis

The *foxo* deletion lines were generated through CRISPR/Cas9 mutagenesis as previously described (Ma et al., 2018). Briefly, two sgRNA plasmids targeting FOXO DNA binding domain were injected into fly embryo. To genotyping G0 flies, single fly was homogenized in 50 μl squashing buffer (10 mM Tris buffer (pH 8.5), 25 mM NaCl, 1 mM EDTA, 200 μg/ml Proteinase K), incubated at 37°C for 30 minutes, then followed by inactivation at 95°C for 10 minutes. Screen primers for *foxo* deletion mutants were: F 5’-GGGGCAGATCCCCGCCCAGC-3’, R 5’-GGGCGATTCGAATAGCAGTGC-3’. The virgin females carrying the deletion were backcrossed into *w^1118^* male flies for five consecutive generations to mitigate background effects.

### Transcriptomic analysis (RNA-Seq)

For transcriptomic analysis on the head tissues of aged flies and *foxo* deletion lines (*foxo^c431^*), forty heads from female flies were dissected and homogenized in a 1.5 ml tube containing 1 ml of Trizol Reagent (Thermo Fisher Scientific, Waltham, MA, USA. Catalog number: 15596026). Three biological replicates were performed for each age and genotype. Total RNA was extracted following manufacturer instruction. TURBO DNA-free kit was used to remove genomic DNA contamination (Thermo Fisher Scientific, Waltham, MA, USA. Catalog number: AM1907). About 1 μg of total RNA was used for sequencing library preparation. PolyA-tailed RNAs were enriched by NEBNext Poly(A) mRNA Magnetic Isolation Module (New England Biolabs (NEB), Ipswich, MA, USA. Catalog number: E7490S). RNA-Seq library was prepared using NEBNext Ultra RNA library Prep Kit for Illumina (NEB, Ipswich, MA, USA. Catalog number: E7420S). The libraries were pooled together in equimolar amounts to a final 2 nM concentration. The normalized libraries were denatured with 0.1 M NaOH (Sigma) and sequenced on the Illumina Miseq or Hiseq 2500 platforms (Single-end, Read length: 100 base pairs) (Illumina, San Diego, CA, USA).

### Chromatin immunoprecipitation sequencing (ChIP-Seq)

Chromatin immunoprecipitation (ChIP) protocol was performed and modified from (Bai et al., 2013). Two biological replicates were collected for each age and genotype. About 200 female flies were first anesthetized with FlyNap (Carolina Biological, Burlington, NC, USA. Catalog number: 173010) and ground into a powder in liquid nitrogen. Crosslinking was performed using 1% paraformaldehyde for 20 minutes followed by glycine quenching. The fly homogenate was washed several times with 1X PBS supplemented with protease inhibitors, and incubated briefly with cold cell lysis buffer (5 mM HEPES pH 7.6, 100 mM NaCl, 1 mM EDTA, 0.5% NP-40). Chromatin was extracted with nuclear lysis buffer (50 mM HEPES pH 7.6, 10 mM EDTA, 0.1% Na-deoxycholate, 0.5% N-lauroylsarcosine), and sheared using Branson digital sonifier 250, using 30%, with 30 seconds on, 30 seconds off for 5 cycles. Chromatin immunoprecipitation was carried out using Protein G SureBeads (Bio-Rad, Hurcules, CA, USA. Catalog number: 1614023). Pre-cleaned chromatin extracts were incubated with anti-FOXO antibody (Bai et al., 2013) and Protein G SureBeads to precipitate FOXO-DNA complexes.

DNA size selection and library prep were done using NEBNext Ultra II DNA library prep kit and indexed using NEBNext multiplex oligos for Illumina (Primer set 1) (NEB, Ipswich, MA, USA. Catalog number: E7645S, E7335S). DNA from either ChIP or input samples was mixed with AMPure XP beads (Beckman Coulter Inc., Brea, CA, USA. Catalog number: A63881) to select for a final library size of 320 bp. Samples were diluted to a final concentration of 2 nM for Illumina sequencing on Illumina HiSeq 3000 (Single-end, Read length: 50 base pairs) (Illumina, San Diego, CA, USA).

### Data processing of RNA-Seq and ChIP-Seq

RNA-Seq reads were first mapped to the reference genome Dm6 with STAR_2.5.3a by default parameter. The read counts for each gene were calculated by HTSeq-0.5.4e. The count files were used as inputs to R package DESeq for normalization. The differential expression genes were computed based on normalized counts from three biological replicates (|log_2_foldchange|>1, adj *p*<0.01).

For ChIP-Seq, raw FASTQ reads were merged using mergePeaks (Homer suite) then uploaded into Galaxy (usegalaxy.org) and checked for quality using FastQC. Files were then run through FASTQ Groomer (https://usegalaxy.org/u/dan/p/fastq) for readability control before mapping reads using Bowtie2 for single-end reads. *D. melanogaster* BDGP Release 6/dm6 was used as the reference genome. BAM output files were converted to SAM using BAM-to-SAM (http://www.htslib.org/doc/samtools.html) and sorted to generate peak images. Peak calling was performed using MACS2. MACS2 FDR (q-value) was set for a peak detection cutoff of 0.05 and did not build the shifting model. The MFOLD for the model was set from 10-50 to detect fold-enrichment. Peak-calling was set to identify peaks 300 bp in length, and no peaks could exceed 10 Kb in size. After MACS2 peak identification, peak regions were expanded 2 kb (1 kb upstream and 1 kb downstream) and assigned to nearby and overlapping genes using BEDTools/intersect (https://bedtools.readthedocs.io/en/latest/content/bedtools-suite.html) with Dm6.16 genome annotation file (UCSC, Santa Cruz, CA, USA). All non-protein coding identified targets were removed from the data set manually based on annotation symbol.

### Venn Diagrams

Venn diagram were created using the Bioinformatics and Evolutionary Genomics Venn calculator at Ugent (http://bioinformatics.psb.ugent.be/webtools/Venn/). For cross species comparisons, gene ID’s were converted to fly ID’s using DIOPT (http://www.flyrnai.org/diopt). Genes that were the best possible match for each ortholog were selected for gene list comparison.

### Quantitative PCR (qPCR)

Quantitative PCR was run on QuantStudio 3 (ThermoFisher Scientific, Waltham, MA USA) with above ChIP and input library samples. PCR reaction was conducted using PowerUp SYBR Green Master Mix (Life Technologies, CA, USA. Catalog number: 4402953). FOXO binding enrichment was determined based on the fold-change between ChIP samples vs. Input samples. The FOXO binding to Actin5C locus was used as a negative control. Two biological and two technical replicates were performed for each age. Primers are listed in Table S2.

### Pathway and gene ontology analysis

Pathway and gene ontology analysis was conducted using Panther (http://www.pantherdb.org/), String (https://string-db.org/) and DAVID (https://david.ncifcrf.gov/). All three methods were used to obtain a more complete picture of shared regulation between datasets. KEGG pathway maps were obtained through KEGG Pathway (http://www.kegg.jp/kegg/pathway.html).

### Motif analysis

Motif analysis was conducted using Homer’s *findMotifsGenome* script (http://homer.ucsd.edu/homer/ngs/peakMotifs.html) to compare peak regions with Dm6.01 FASTA data from UCSC.

### List of raw datasets used

ChIP-Seq datasets: GSE62580 (*Drosophila* aging fat body), GSE81100 (*Drosophila* aging head tissue), GSE44686 (*Drosophila chico* heterozygotes FOXO ChIP), GSE15567 (Encode *C. elegans Daf-16* ChIP)

### Immunofluorescent staining

Flies were anesthetized with FlyNap and dissected in 1X PBS. Fly tissues (muscle or fat body) were then fixed in 4% paraformaldehyde for 20 minutes at room temperature. Tissue was washed in 1X PBST (0.1% Triton X) and blocked with 5% normal goat serum (NGS) for 1 hour at room temperature. Fly tissues were stained with anti-FOXO antibody in 1X PBST at a dilution of 1:1000 for 16 hours at 4°C on a rotator. Tissues were placed in secondary anti-body goat-anti-rabbit conjugate Alexa Fluor 488 (Jackson ImmunoResearch Laboratories Inc., West Grove, PA, USA) at a Dilution of 1:250 and kept in the dark at room temperature for 2 hours. The nucleus was stained using SlowFade with DAPI. Images were captured using an epifluorescence-equipped BX51WI microscope (Olympus, Waltham, MA, USA). Image deconvolution was conducted using CellSens software (Olympus, Waltham, MA, USA), and compiled using ImageJ Fiji.

### Statistical analysis

GraphPad Prism (GraphPad Software, La Jolla, CA, USA) was used for statistical analysis and to generate Boxplot. To compare the mean value of treatment groups versus that of control, either student t-test or one-way ANOVA was performed using Dunnett’s test for multiple comparison.

## Acknowledgments

We thank Bloomington Drosophila Stock Center for fly stocks. We thank Michael Baker and DNA Facility at Iowa State University (ISU) for help with RNA-Seq analysis, Usha Muppirala and Andrew Severin from ISU Genome Informatics Facility (GIF) for assistance with bioinformatics. We thank Christian Riedel for providing *C. elegans Daf-16* ChIP-Seq data. This work was supported by NIH/NIA R00 AG048016 to HB, AFAR Research Grants for Junior Faculty to HB, and National Program on Key Basic Research Project of China 2016YFA0501900 to NL.

## Author’s Contributions Statement

Conceived and designed the experiments: AB XW NL HB. Performed the experiments: AB XW. Analyzed the data: AB XW MT NL HB. Wrote the paper: AB XW NL HB. All authors reviewed and approved the manuscript.

## Availability of data and materials

The raw data files of sequencing experiments have been deposited in the NCBI Gene Expression Omnibus. The accession number for RNA-Seq data is GSE122470 (https://www.ncbi.nlm.nih.gov/geo/query/acc.cgi?acc=GSE122470). The accession number for ChIP-Seq data is GSE121102 (https://www.ncbi.nlm.nih.gov/geo/query/acc.cgi?acc=GSE121102).

## Supporting information

**Figure S1.**
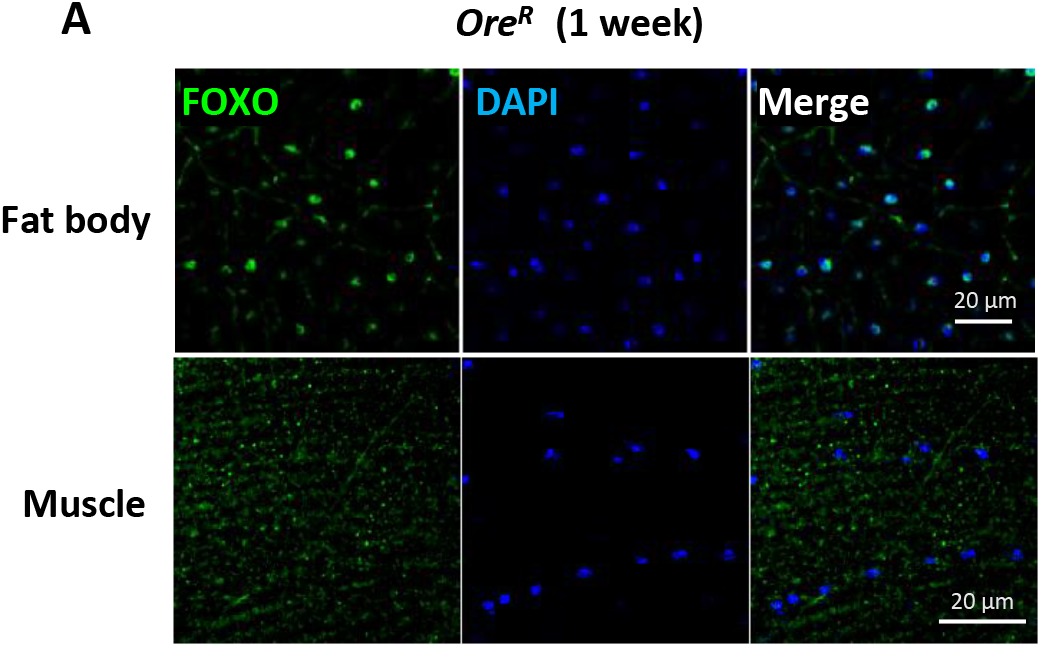
Abdominal fat body and flight muscle of wild-type flies (*Ore^R^*) stained with anti-FOXO at young (2 weeks) and old age (5 weeks). Scale bar: 20 μm.

**Figure S2.**
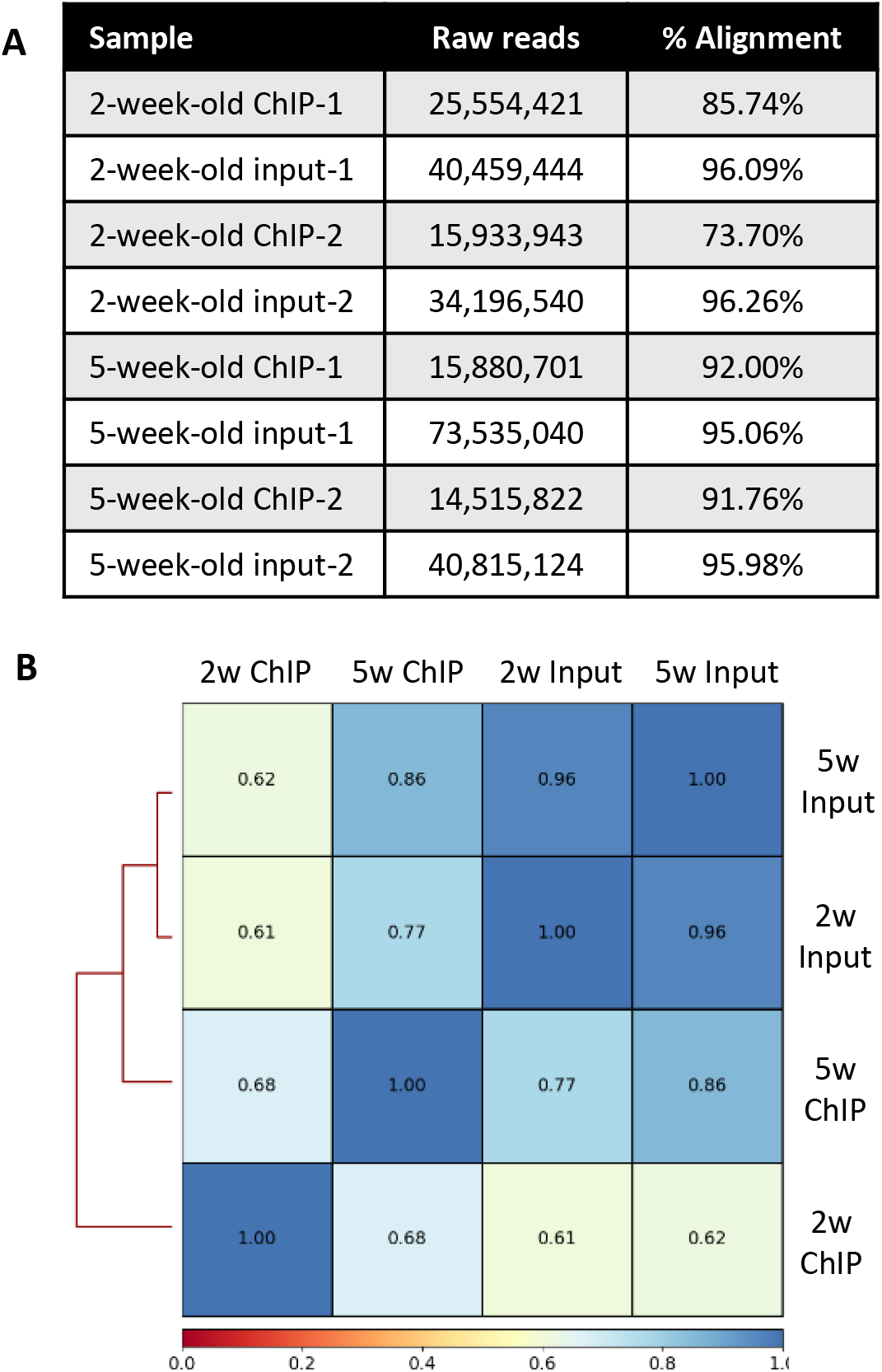
**A)** The total number of raw reads and Bowtie alignment percentage for individual ChIP-Seq sample. **B)** Plot correlation matrix showing the overall correlation among young and old ChIP and input samples.

**Figure S3.**
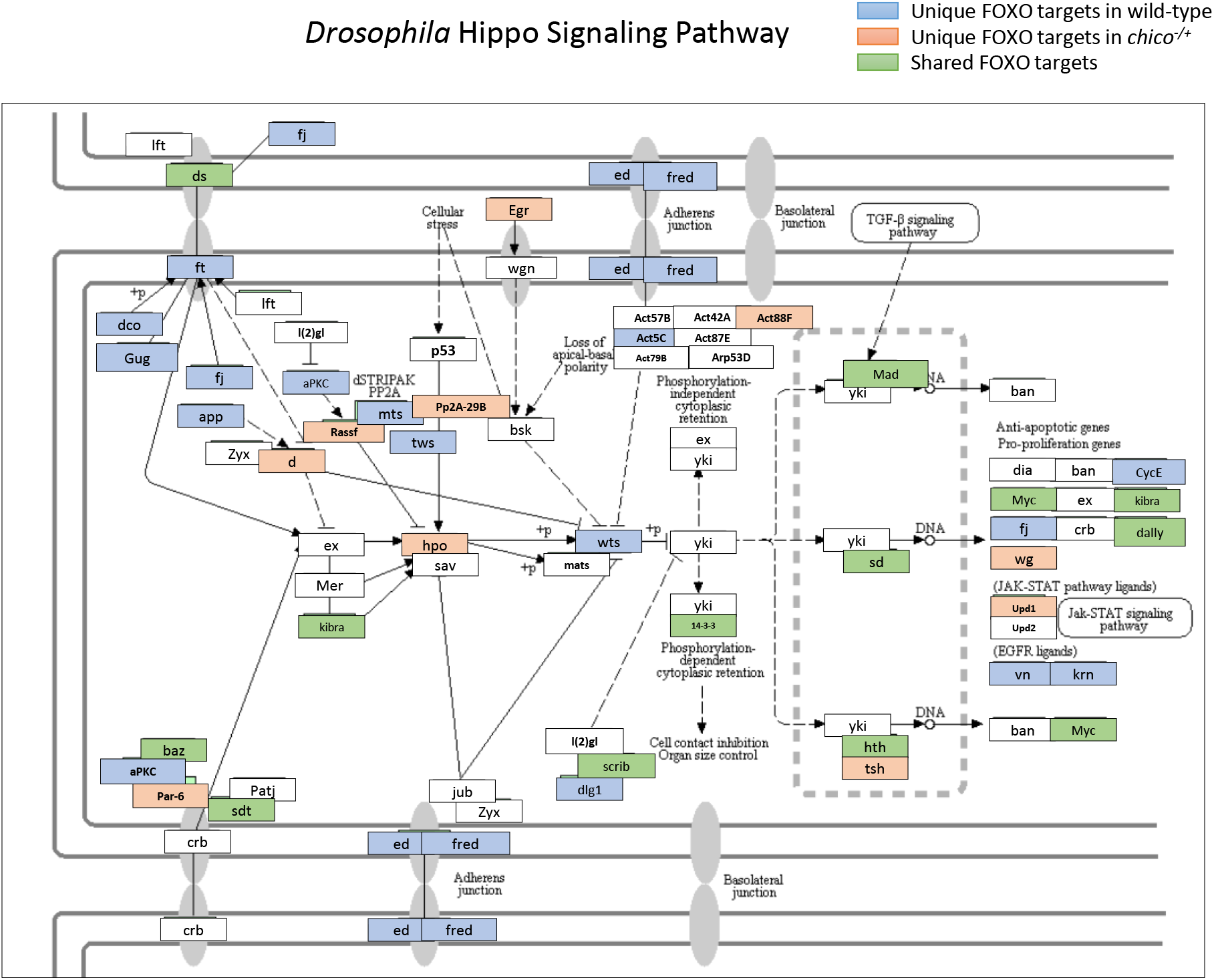
FOXO target genes in Hippo signaling pathway. Unique FOXO targets in wild-type flies (*yw^R^*) are highlighted in blue. Unique FOXO targets in *chico* mutants are highlighted in orange. Shared targets are highlighted in green.

**Figure S4.**
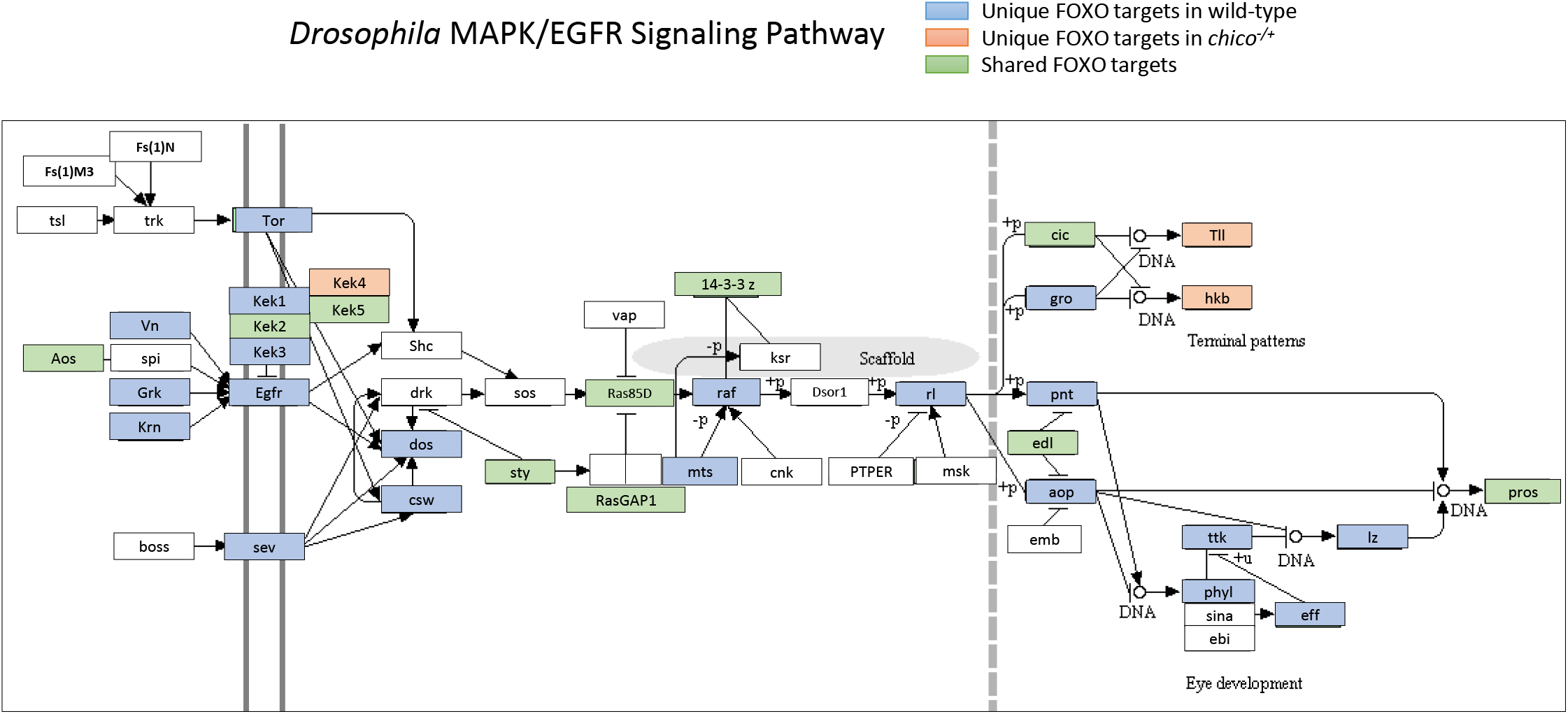
FOXO target genes in MAPK/EGFR signaling pathway. Unique FOXO targets in wild-type flies (*yw^R^*) are highlighted in blue. Unique FOXO targets in *chico* mutants are highlighted in orange. Shared targets are highlighted in green.

**Table S1:**

Lists of peaks, target genes, and GO terms

**Table S2:**
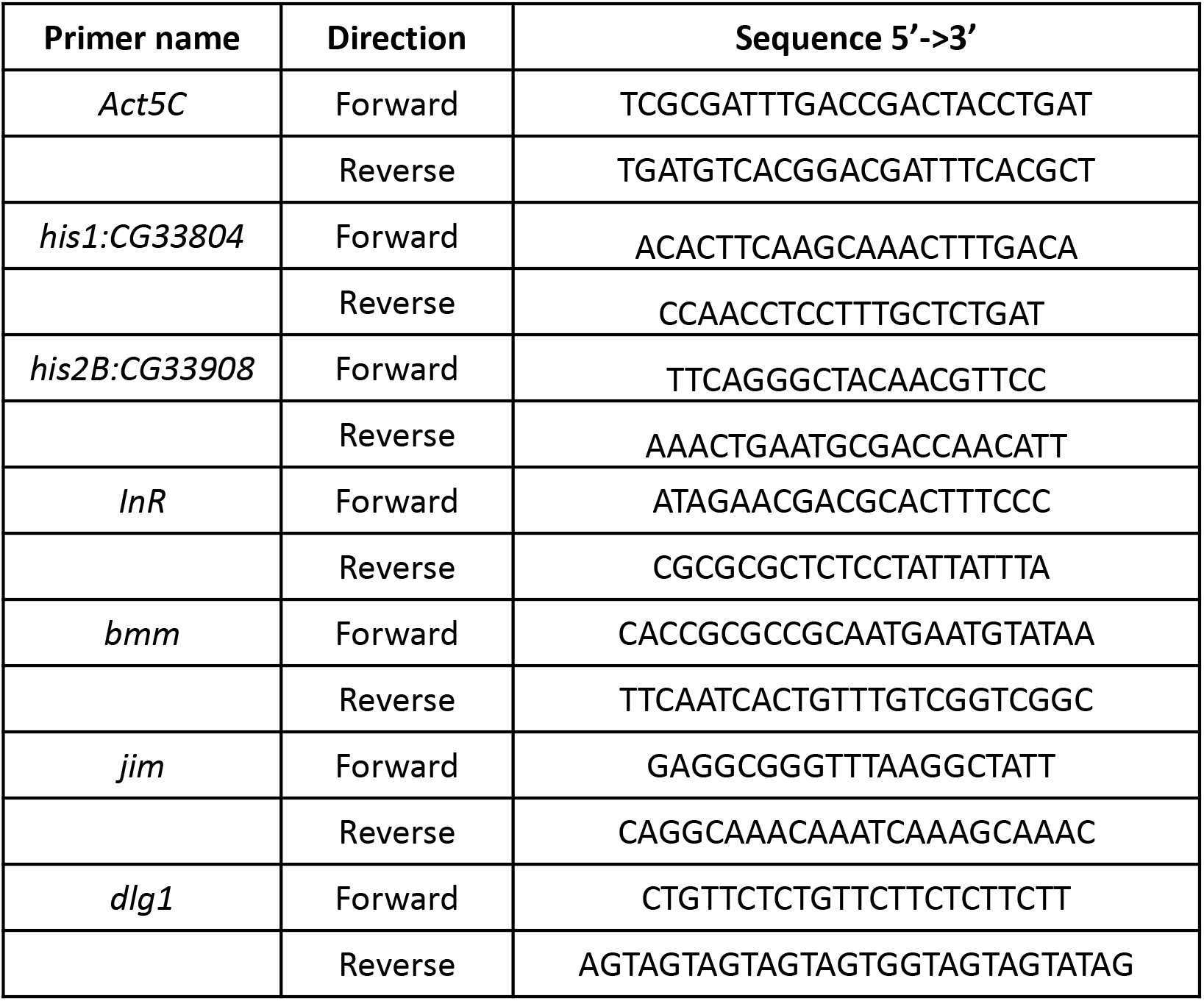
Lists of primers used in qPCR analysis

